# A dual-function regulatory element couples ParB expression and DNA substrate specificity

**DOI:** 10.64898/2026.07.07.737113

**Authors:** Rodrigo A. Arana-Garrido, Daniel Lee Joo, Huda Al-Zuhairi, John D. Norcross, Ariana N. Nielsen, HyeongJun Kim, Paola E. Mera

**Author notes:** Equal contributors (Rodrigo A. Arana-Garrido, Daniel Lee Joo).

## Abstract

Chromosome segregation is essential for cell survival. Most bacteria encode the chromosome partitioning ParABS system. Although even small changes in ParA or ParB levels disrupt genome maintenance, the mechanisms that control their abundance have remained unresolved. Using *Caulobacter crescentus*, we provide the first mechanistic evidence that ParB levels are regulated post-transcriptionally. Through single-nucleotide substitutions and compensatory mutation analyses, we identify an mRNA secondary structure at the 5′ end of the *parB* transcript that enhances ParB cellular abundance. Loss of this regulatory mechanism sensitizes cells to modest increases in ParA levels, causing cell death. Additionally, we demonstrate that ParB substrate specificity is not determined solely by the central helix–turn–helix domain that interacts with *parS* but is unexpectedly modulated by the N-terminal domain. Together, our findings reveal an uncharacterized layer of chromosome segregation control and highlight post-transcriptional regulation of ParB abundance as a potential mechanism for maintaining the precise ParA–ParB balance required for bacterial viability.

## INTRODUCTION

The faithful inheritance of chromosomes is essential for the survival and proliferation of bacteria. To ensure proper genome inheritance, approximately 70% of bacterial species rely on the conserved partitioning system ParABS [1]. This system is composed of two proteins, ParA and ParB, and a centromere-like chromosomal locus (*parS*). The essential role of ParABS in chromosomal maintenance has been demonstrated across a wide range of bacteria, including asymmetrically dividing, spore-forming, predatory, multipartite genome-containing, and pathogenic species [2–7]. Perturbations in the ratio between cellular ParA and ParB levels compromise the important processes in bacteria, such as chromosome partitioning and cell division, and can result in cell death [8–10]. Despite this sensitivity, the mechanisms by which bacteria maintain the appropriate balance between these proteins remain poorly understood.

ParA contains a P-loop ATPase motif and functions as the motor-like component of the partitioning system. Upon adenosine triphosphate (ATP) binding, ParA dimerizes, enabling its nonspecific association with the nucleoid [11–15]. ParB, bound at *parS* sites, stimulates ParA ATP hydrolysis, leading to the release of ParA monomers from DNA [16, 17] and generating a dynamic gradient of ParA [18–20]. The ability of ParA to cycle between ATP-bound and ADP-bound states is central to its function in driving directional chromosome segregation. Aside from its canonical role in partitioning, ParA has also been implicated in coordinating DNA replication initiation and in organizing chromosome architecture [21–23]. Notably, these additional functions appear to depend on the nucleotide-bound state of ParA, highlighting the importance of its biochemical cycle in integrating multiple chromosome-associated processes.

ParB is a distinctive DNA-binding protein that functions as a nucleotide-dependent molecular switch, utilizing cytidine triphosphate (CTP) rather than the more commonly employed ATP [24–26]. It exhibits high affinity for centromere-like *parS* sequences, which are typically found proximal to the origin of replication (*oriC*) [1, 27, 28]. Upon binding to *parS*, CTP association induces a conformational transition in ParB that promotes the formation of a sliding clamp, enabling it to spread along adjacent DNA [26, 29–32], in a manner reminiscent of the eukaryotic processivity factor proliferating cell nuclear antigen (PCNA) [33]. This spreading is thought to be central to the formation of a higher-order nucleoprotein complex required for efficient partitioning [26, 34]. Beyond its direct role in *parS*-based segregation, ParB serves as a key hub for chromosome organization by recruiting the structural maintenance of chromosomes (SMC) complex [35, 36], interacting with polar chromosomal anchoring factors [37–40], and coordinating the spatial regulation of division inhibitors [9]. ParB proteins from diverse bacterial species share three common structural domains: (1) ParB proteins dimerize through the C-terminal domain (CTD) [30, 33, 41, 42]. (2) The central DNA-binding domain (CDBD) features a specific *parS* DNA sequence recognition and binding via a helix-turn-helix motif [24, 29, 30, 43]. (3) A CTP-binding region resides at the N-terminal domain (NTD). Additionally, the consensus sequence (LGXGL) at NTD is responsible for the interaction between ParA and ParB [11, 44, 45]. Given the essential and multifaceted roles of ParB, even modest perturbations in its cellular levels can be detrimental or lethal to the cell [6–8, 46, 47].

Perturbations in the balance between ParA–ATP dimers and ParA–ADP monomers impact chromosome segregation and result in pleiotropic defects, reflecting the broader influence of ParA’s nucleotide-bound state on replication initiation and chromosome organization [6, 21, 48, 49]. Because ParB stimulates the hydrolysis of ParA–ATP dimers into monomers, changes in ParB levels directly alter the distribution of ParA nucleotide states [10, 50]. Consistent with this coupling, co-upregulation of ParA and ParB can often rescue the observed defects [9, 47, 49]. While *parA* and *parB* are encoded within the same operon, little is known about how their expression is regulated. Reports of transcriptional autoregulation by ParA and ParB largely derive from plasmid-encoded systems and are not well established for chromosomal ParABS systems [51–54]. Thus, a fundamental unresolved question is how bacteria maintain optimal ParA–ParB levels to ensure proper chromosomal maintenance.

One of the best-characterized chromosomal ParABS systems is found in *Caulobacter crescentus*. Following the onset of chromosome replication, the duplicated *parS* loci, decorated with ParB, are segregated from the stalked pole to the opposite new pole by a ParA gradient that extends from the new pole toward the stalked pole [20, 55, 56]. Mechanistic features described in other bacteria have also been demonstrated in *C. crescentus*, including the nucleotide cycles of ParA–ATP and ParB–CTP, CTP-dependent spreading of ParB from parS, recruitment of the SMC complex to *parS* by ParB, and the connection between ParA and replication initiation [26, 30, 48, 55–57]. *C. crescentus* provides a powerful model to study the chromosomal ParABS system because it divides asymmetrically and initiates replication once per cell cycle [58], allowing clear assessment of segregation fidelity and directionality [58].

Using *C. crescentus*, we uncover a previously unrecognized layer of regulation governing ParB levels at the post-transcriptional level. We demonstrate that this regulation is mediated by a secondary structure formed within the *parB* gene at the 5’-end. Strikingly, we show that the first 10 amino acids on the *C. crescentus* ParB determines its specificity for *parS*, in addition to the helix-turn-helix motif at the central DNA-binding domain (CDBD), expanding current models of ParB–DNA interaction.

Together, these findings provide the first mechanistic evidence that cells actively tune ParB abundance.

## RESULTS

### Two sized ParB proteins interchangeably used

While examining ParB’s impact on cell viability, we noticed that *parB* in *C. crescentus* has two potential start codons (ATG) separated by 30 nucleotides (Figure 1A). Analysis of the literature revealed that these two start codons have been used interchangeably. Here, we refer to the protein version starting at the first methionine as "full-length ParB (FL)" and the version starting from the second start codon as "delta-10 ParB (Δ10)". To determine potential differences associated with initiation at either start codon, we analyzed the effects of increased ParB expression on cell viability. To increase the levels of ParB, we constructed *parB* merodiploid strains by incorporating a second *parB* copy (either full-length or delta-10) expressed from the chromosomal xylose-inducible promoter (Pxyl), leaving the native *parB* gene intact (Figure 1B). Using strains expressing *parB* full-length, delta-10, and empty vector (EV) as control, we examined cell morphology and viability. Surprisingly, our data revealed that the phenotypes were significantly different depending on which version of *parB* was being expressed. Upon 6 h of induction, cells expressing the Pxyl-full-length *parB* displayed filamentous morphology and significant loss of viability (Figure 1CD). However, cells expressing Pxyl-delta-10 *parB* displayed similar cell morphology and viability distributions to the empty vector control.

**Figure 1.**
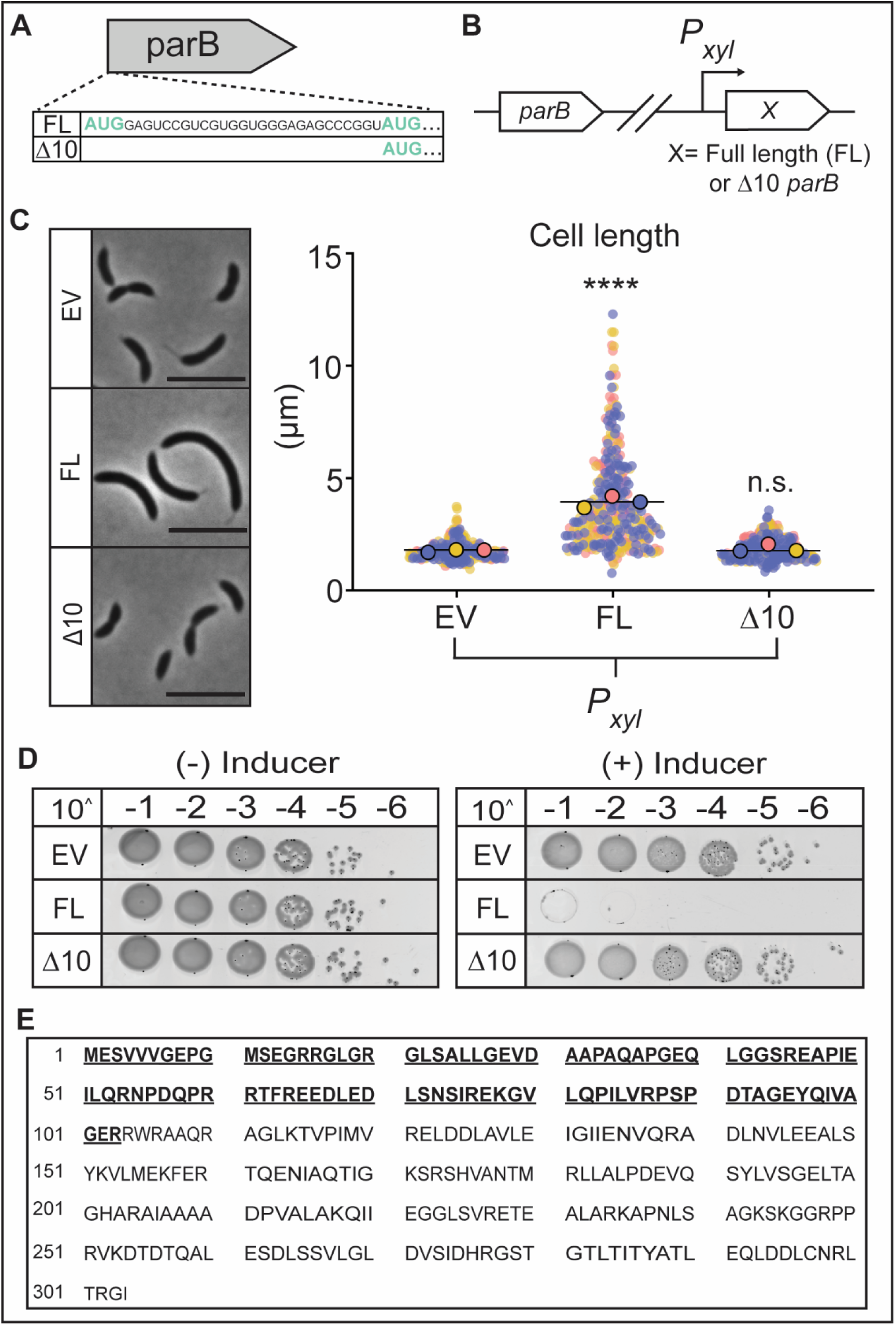
Cells are sensitive to overexpression of full-length but not of Δ10 ParB. **A.** Schematic of the *parB* gene showing the first 33 nucleotides. Highlighted in teal are the two AUG codons present in the *parB* gene. The first codon encodes for Full-length ParB (FL), and the second codon encodes for Δ10 ParB (Δ10). **B.** Schematic of FL and Δ10 ParB inducible strains. **C.** (Left) Phase contrast images of FL and Δ10 inducible strains after 6h of 0.1% xylose induction, grown in rich media (PYE) to exponential phase, scale 5 µm. (Right) Cell length quantification of mixed population. Small points represent data points from three independent replicates, large dots represent median values (blue, pink, yellow). The horizontal lines represent the mean of three median values. One-way ANOVA was performed using population mean values (N=3) to compare values for each measurement. p(<0.0001) ****, n.s. (non-significant). The analyses were blinded n=∼400 cells. **D.** Colony-forming units showing viability of inducible strains in comparison to empty vector (EV). EV and Δ10 show similar growth, while FL shows a viability defect. **E.** Amino acid sequence of ParB. The amino acids validated by Liquid Chromatography–Mass Spectrometry (LC-MS) are bolded and underlined.

Given the known sensitivity to increased ParB levels in various bacterial species [6–8, 46, 47], the defect observed upon FL *parB* overexpression suggested that the full-length form is the one produced in *C. crescentus* cells. To further examine this possibility, we performed mass spectrometry analysis on cell-free extracts from wild-type cells (Figure 1E). Using a gel slice from an SDS–PAGE corresponding to the predicted molecular weight of both ParB sizes, the peptides detected by mass spectrometry matched across the entire N-terminus, confirming that full-length ParB is synthesized in wild-type *C. crescentus* cells. Production of the full-length ParB is also consistent with ribosome profiling data showing that *parB* translation initiates at the first start codon [59]. Beyond identifying which start codon is used in *C. crescentus*, these results suggested a previously unrecognized layer of regulation of ParB protein levels.

### The 5’ end of parB determines levels of ParB protein

To further investigate the mechanism of a potential regulation of ParB that results in the distinct phenotypic differences observed upon expression of full-length or Δ10 *parB*, we examined how each protein variant accumulated over time. Because we used a *parB* merodiploid strain to increase the expression levels of either full-length or Δ10 *parB*, we needed a way to distinguish protein levels originating from the inducible promoter versus those from the native *parB* promoter. Attempts to generate a ParB depletion strain, in which *parB* with either start codon expression was placed under an inducible promoter and the native locus deleted, were unsuccessful. To overcome this limitation, we used *parB* merodiploid strains in which the native *parB* gene was tagged with a fluorescent protein, thereby increasing the molecular weight of the ParB expressed from the native promoter (CFP-ParB: 58.59 kDa vs. ParB: 32.81 kDa) and enabling clear differentiation of molecular weights by western blot. We confirmed that this strain, just like WT untagged *parB*, responds with the same pattern of morphology defect and loss of viability when overexpressing full-length or Δ10 *parB* from Pxyl (Supplementary Figure 1). To quantify the accumulation of full-length and Δ10 ParB proteins, we performed western blot analysis following induction of the xylose promoter, using empty vector (EV) as a negative control. Over a 2-hour period, xylose was added to the cultures, and samples were collected at 30-minute intervals. As expected, the EV control showed no detectable untagged ParB expression throughout the experiment (Figure 2A). In contrast, both full-length and Δ10 ParB constructs exhibited progressive accumulation of their respective proteins. Notably, while both ParB versions were expressed, the Δ10 ParB accumulated to significantly lower levels (∼10%) than the full-length ParB. These findings revealed two important insights: first, that the higher accumulation of full-length ParB compared to Δ10 is likely responsible for the observed morphological defects and loss of viability; and second, more importantly, that the 5′ end of *parB* or the N-terminus of ParB plays a key role in regulating ParB cellular accumulation.

**Figure 2.**
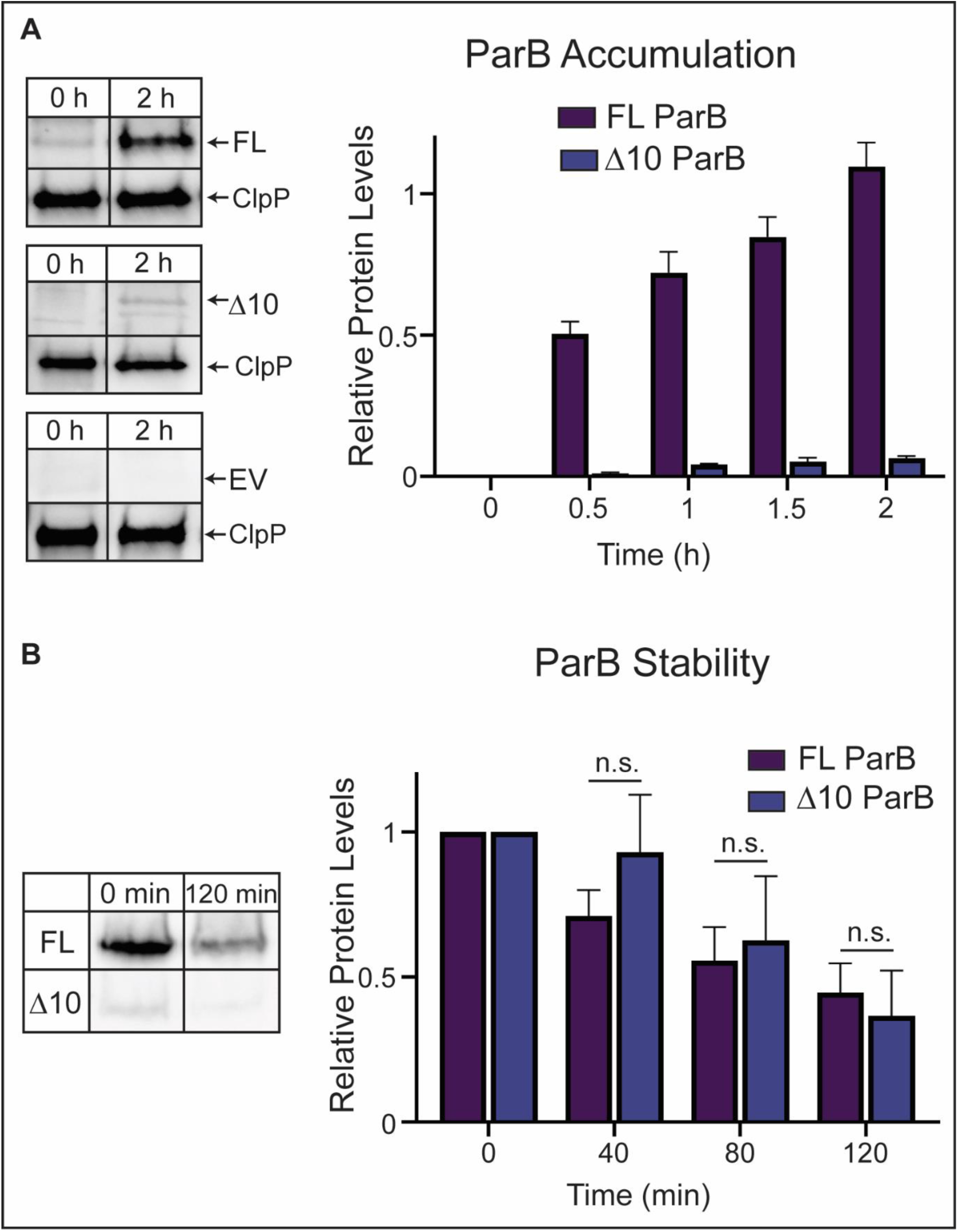
The 5’ end of parB determines levels of ParB protein. **A.** Quantification of ParB protein accumulation over time by western blot analysis. Cells were induced with 0.1% xylose, and samples were collected every 30 minutes. Data were obtained from triplicate independent samples. Band intensity was measured and normalized onto loading control, ClpP. (See also Supplemental Figure 3A.) **B.** Quantification of ParB protein degradation over time by western blot analysis. Cells were induced for 1 hour before chloramphenicol treatment (10 µg/ml) to halt protein synthesis. Samples were then collected every 40 minutes. Data were obtained from triplicate independent samples. Band intensity was measured and normalized onto band intensity recorded at time = 0 after addition of chloramphenicol. Two-way ANOVA was performed to compare values at each time point. (See also Supplemental Figure 3B.)

### ParB levels are regulated at the post-transcriptional level

We first examined whether proteolysis of full-length versus Δ10 ParB contributed to the observed differences in protein accumulation. To test this possibility, we analyzed the stability of both full-length ParB and the Δ10 variant inside cells over time. Using exponential-phase cultures following induction, we stopped protein synthesis by adding a low concentration of the ribosomal inhibitor chloramphenicol [60]. Samples were collected at 40-minute intervals, and protein abundance was monitored over time using western blots (Figure 2B). These data showed that the stability of full-length and Δ10 ParB is similar, eliminating proteolysis as the source of the differential accumulation of ParB.

We next examined whether *parB* was regulated post-transcriptionally. We reasoned that if the regulation of ParB levels is post-transcriptional, the nucleotide sequence, and not the amino acid sequence, at the 5′ end of *parB* would be responsible for modulating expression levels. To test this hypothesis, we engineered 5 silent mutations within the first 30 base pairs of the full-length *parB* that altered the nucleotide sequence without changing the encoded sequence of the 10 amino acids, referred here as *parB*-silent5 or S5 (Figure 3A). Notably, the western blot data revealed that by simply changing 5 bases without impacting the amino acid sequence, the levels of accumulation of the ParB-silent5 protein were significantly reduced compared to the accumulation of full-length ParB both expressed from Pxyl (Figure 3B). Consistent with these reduced accumulation levels, expression of *parB*-silent5 resulted in cells with decreased filamentation (Figure 3C) and higher viability (Figure 3D) than cells expressing unaltered FL ParB. These results suggested that ParB accumulation levels are regulated post-transcriptionally.

**Figure 3.**
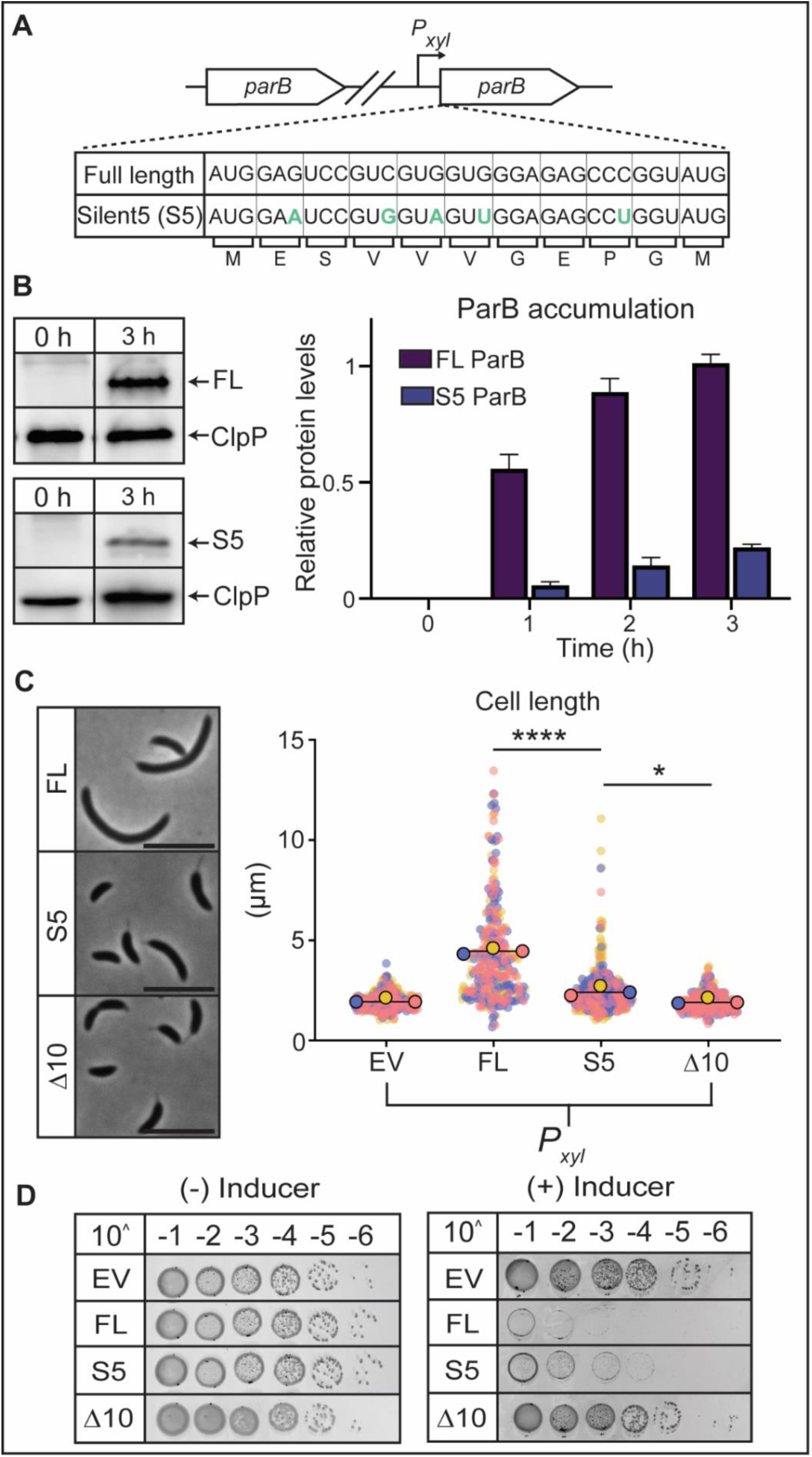
ParB levels are regulated at the post-transcriptional level. **A.** Schematic of ParB merodiploid strains. Silent 5 (S5) ParB contains 5 nucleotide changes on the *parB* gene highlighted in teal. **B.** Quantification of ParB protein accumulation over time by western blot analysis. Cells were induced with 0.1% xylose, and samples were collected every hour. Data obtained from triplicate independent samples. Band intensity was measured and normalized to the loading control, ClpP. (See also Supplemental Figure 3C.) **C.** (Left) Phase contrast images of FL, Δ10, and S5 ParB inducible strains after 6h of induction, grown in rich media (PYE) to exponential phase, scale 5 µm. (Right) Cell length quantification of mixed population. Small points represent data points from three independent replicates, large dots represent median values (blue, pink, yellow). The horizontal lines represent the mean of three median values. One-way ANOVA was performed using population mean values (N=3) to compare values for each measurement. p(<0.0001) ****, p(<0.05) *, n.s. (non-significant). All samples were blinded n=∼500 cells. **D.** Colony-forming units showing viability of inducible strains in comparison to empty vector EV. EV and Δ10 show similar growth, while FL shows a viability defect. S5 shows a milder viability defect to FL.

### Post-transcriptional regulation involves a regulatory mRNA secondary structure

Having established that changes within the first 30 base pairs are sufficient to alter ParB protein accumulation levels, we posit that post-transcriptional regulation of *parB* is mediated by a regulatory RNA secondary structure formed within the 5’ region of the mRNA. To guide our analysis, we used RNA-Fold [61] to model potential structures formed by the 30 base pairs. RNA-Fold predicted the formation of a stable stem-loop structure by the native 30 base pairs at the 5’-end of *parB* (Figure 4A). Notably, the mutations in *parB*-silent5, which were designed prior to knowledge of the RNA-Fold-predicted secondary structure, destabilized the native stem loop and led to the formation of three large loops (Figure 4B). The significant changes in the secondary structure at the 5’-end in *parB*-silent5 were consistent with our post-transcriptional hypothesis explaining the reduced expression levels of *parB*-silent5 compared to the unaltered full-length ParB (Figure 3B).

**Figure 4.**
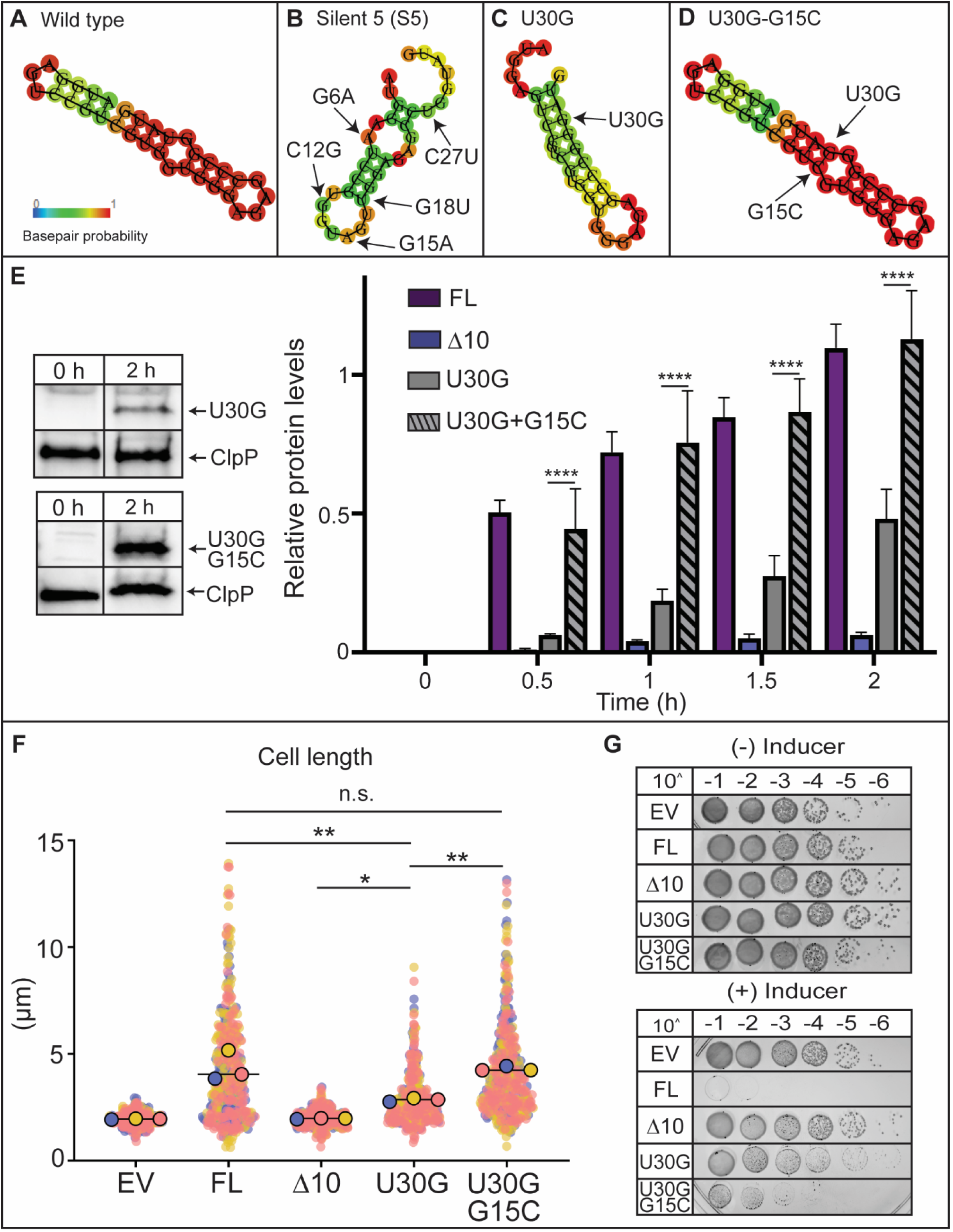
Post-transcriptional regulation involves a regulatory RNA secondary structure. **A-D.** Secondary structure predictions of the first 33 nucleotides of the *parB* transcript by RNAfold. Probability of base pair formation is displayed in a color gradient based on the stability of such base pair. **E.** (Left) Western blot analysis of U30G and U30G G15C. Cells were induced with 0.1% xylose, and samples were collected every 30 minutes. Quantification of ParB protein accumulation over time by western blot analysis. Data were obtained from triplicate independent samples. Band intensity was measured and normalized onto loading control, ClpP. Two-way ANOVA was performed to compare values at each time point. p(<0.0001) ****. (See also Supplemental Figure 3D.) **F.** Cell length quantification of mixed population. Small points represent data points from three independent replicates, large dots represent median values (blue, pink, yellow). The horizontal lines represent the mean of three median values. One-way ANOVA was performed using population mean values (N=3) to compare values for each measurement. p(<0.01)**, p(<0.05)*n.s. (non-significant). All samples were blinded n=∼300 cells. **G.** Colony-forming units showing viability of inducible strains in comparison to empty vector (EV). EV and Δ10 show similar growth while FL and U30G-G15C show a viability defect. U30G shows a milder viability defect to FL.

The *parB*-silent5 allele introduced two rare codons, raising the possibility that reduced translational efficiency rather than secondary structure disruption contributed to decreased accumulation (Supplemental Table 2). To eliminate the possibility of a codon-dependent change in ParB levels, we used RNA-fold to design new mutations that directly disrupt the predicted stable stem–loop without affecting codon usage. We identified a single base substitution at nucleotide position 30 (uracil to guanine, U30G) predicted to destabilize the secondary structure at the 5′ end of the *parB* transcript (Figure 4C). We confirmed that the native codon GGU and the substituted codon GGG exhibit equivalent codon-usage frequencies in *C. crescentus* (Supplemental Table 2), thereby eliminating potential differences in translational efficiency. Remarkably, introducing this single base substitution at position 30 resulted in a decrease in ParB accumulation levels compared to the unaltered FL ParB (Figure 4E), consistent with the loss of stability of the mutant stem loop. In line with the reduced levels of ParB accumulation, the phenotypes of cells expressing the substitution Uracil-30-Guanine also resembled those of the *parB*-silent5 (Figure 4FG).

To validate the presence of this secondary structure and its regulatory role, we took advantage of the base-pairing properties of RNA and engineered a compensatory mutation. Using RNA-Fold, we confirmed that substituting guanine at position 15 with cytosine (guanine to cytosine, G15C) restores the base-pair complementation of uracil-30-guanine, reestablishing the formation of the original stable stem-loop (Figure 4D). Our analysis showed that the combined U30G and G15C substitutions restored ParB protein accumulation to levels equivalent to unaltered FL ParB, leading to filamentous cell morphology and loss of viability (Figure 4E-G), confirming the regulatory role of the predicted secondary mRNA structure. Together, our findings show that ParB levels are regulated post-transcriptionally through a stable secondary structure within the 5’-end of the *parB* transcript.

### Evidence for post-transcriptional regulation at the native parB chromosomal locus

Because *parB* is essential in *C. crescentus*, attempts to introduce modifications directly at the native locus frequently yield clones that have reverted to the wild-type sequence, likely due to selective pressure against changes in ParB expression during strain construction. We therefore initially relied on merodiploid strains where all the modifications at the 5′-end were engineered in the *parB* copy expressed from the xylose-inducible promoter. However, *parB* has an independent transcription start site within the *gidAB–parAB* operon [62], suggesting that its native regulatory context could influence its expression. We therefore examined whether the post-transcriptional regulation of *parB* observed in merodiploid strains also occurs when *parB* is expressed from its native chromosomal locus, and whether this regulation is impacted by the native promoter or by the 5′ untranslated region (5’UTR). We successfully introduced a chromosomal nucleotide modification at the third position (guanine to adenine, G3A). RNA-fold predictions indicated that this substitution altered the RNA secondary structure by generating a large, stable loop absent in the original structure (Figure 5A). Consistent with our findings in the merodiploid strain, the chromosomal G3A mutant showed a clear reduction in ParB protein accumulation relative to wild-type levels (Figure 5B), supporting the model in which post-transcriptional regulation of *parB* occurs at its native chromosomal locus.

**Figure 5.**
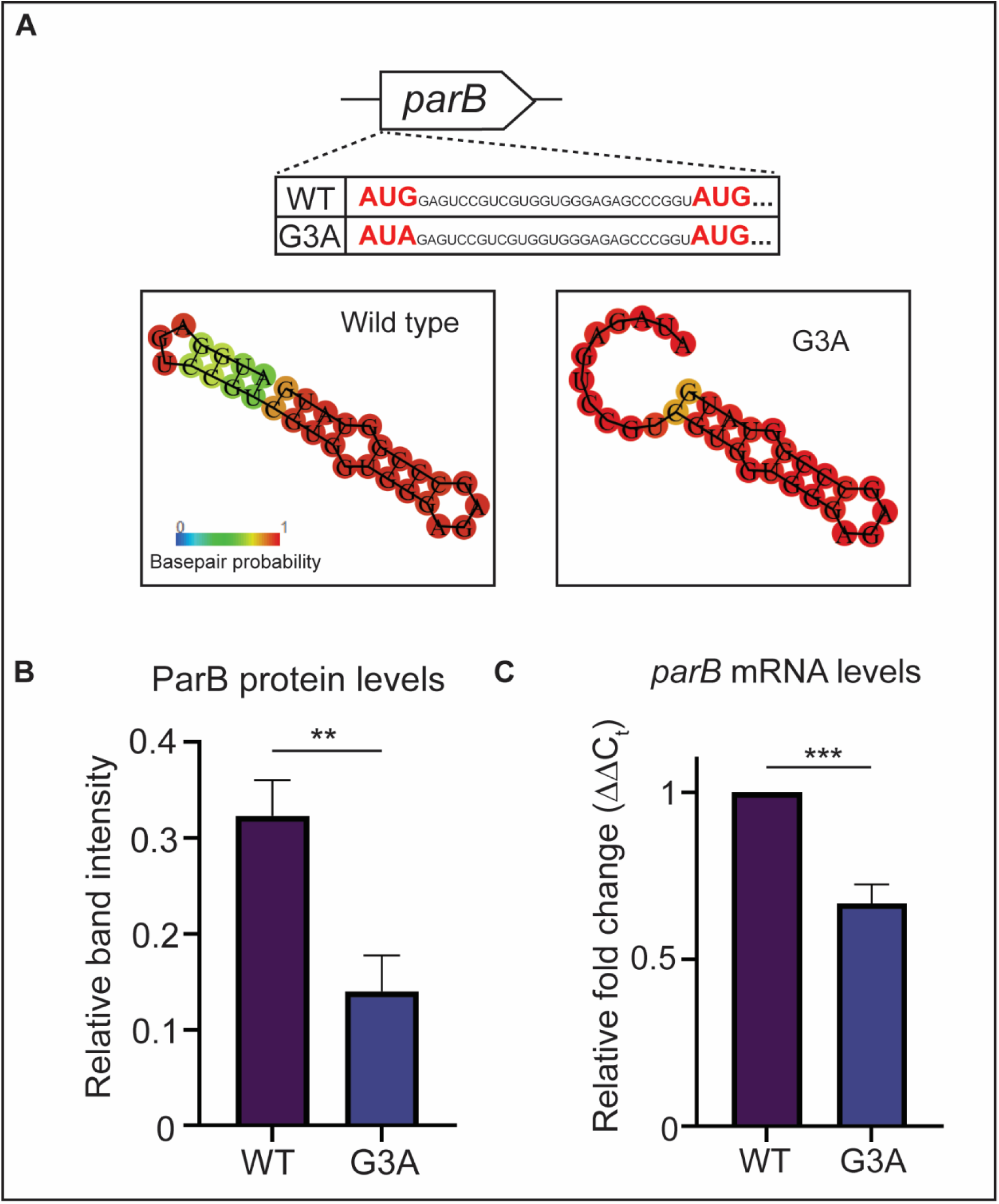
Post-transcriptional regulation confirmed from *parB*’s native chromosomal locus. **A.** (Upper) Schematic of the native *parB* gene. G3A has a single nucleotide mutation in the third nucleotide of *parB* (G3A). (Lower) Secondary structure predictions of the first 33 nucleotides of the parB transcript by RNAfold. Probability of base pair formation is displayed in a color gradient based on the stability of such base pair. **B.** Quantification of ParB protein expression from the native promoter by western blot analysis. Data were obtained from triplicate independent samples. Band intensity was measured and normalized onto loading control, ClpP. Unpaired t-test was performed to compare band intensity values. p(<0.01)**. C. RT-qPCR analysis of *parB*. Cultures were grown to log phase, and RNA was harvested to measure gene expression. Gene expression was normalized relative to the expression of *ruvA*. Unpaired t-test was performed to compare mRNA abundance values. p(<0.001)***.

Because the G3A mutation was present on the chromosome without additional copies of *parB*, we could directly compare transcript levels to those of wild-type cells. Using reverse transcription quantitative PCR (RT–qPCR) to quantify *parB* transcript levels, we found that the G3A chromosomal mutation within the *parB* coding sequence resulted in a significant reduction in transcript abundance compared to wild-type cells (Figure 5C). Collectively, these findings support a model in which *parB* expression is regulated at the post-transcriptional level through a secondary structure located at the 5′ end of the transcript, with disruption of this structure leading to reduced mRNA accumulation and, consequently, decreased ParB protein levels.

### Δ10 ParB-only strains are highly susceptible to ParA fluctuations

Aside from altering the predicted secondary structure of the mRNA, the chromosomal G3A substitution disrupted the first methionine codon by converting it to isoleucine, redirecting translation to the 11^th^ AUG codon and thus producing Δ10 ParB from the native locus. Our ability to successfully construct this strain demonstrated that *parB* is synthesized from this mutant strain and that the resulting ParB protein is sufficient to support cell viability. These results raised the question of why post-transcriptional regulation is required if expression of the truncated ParB at low levels is sufficient for survival. We hypothesized that the importance of post-transcriptional regulation of *parB* may be linked to ParA, as proper ParB levels are critical for maintaining the correct balance between ParA dimers and monomers in the cell [10, 50]. To test this hypothesis, we constructed *parA* merodiploid strains to examine the ability of cells carrying the G3A mutation at the native *parB* locus to tolerate modest increases in ParA levels (Figure 6A). In contrast to wild-type cells, the G3A mutant strain exhibited a pronounced increase in cell length within 3h of *parA* induction (Figure 6B). While wild-type *parB* cells readily accommodated subtle increases in ParA levels [63], the same ParA increases result in G3A cells loss of viability (Figure 6C). Together, these results demonstrate that loss of *parB* post-transcriptional regulation sensitizes cells to fluctuations in ParA levels, which can result in loss of viability, thereby highlighting the critical role of this regulatory mechanism.

**Figure 6.**
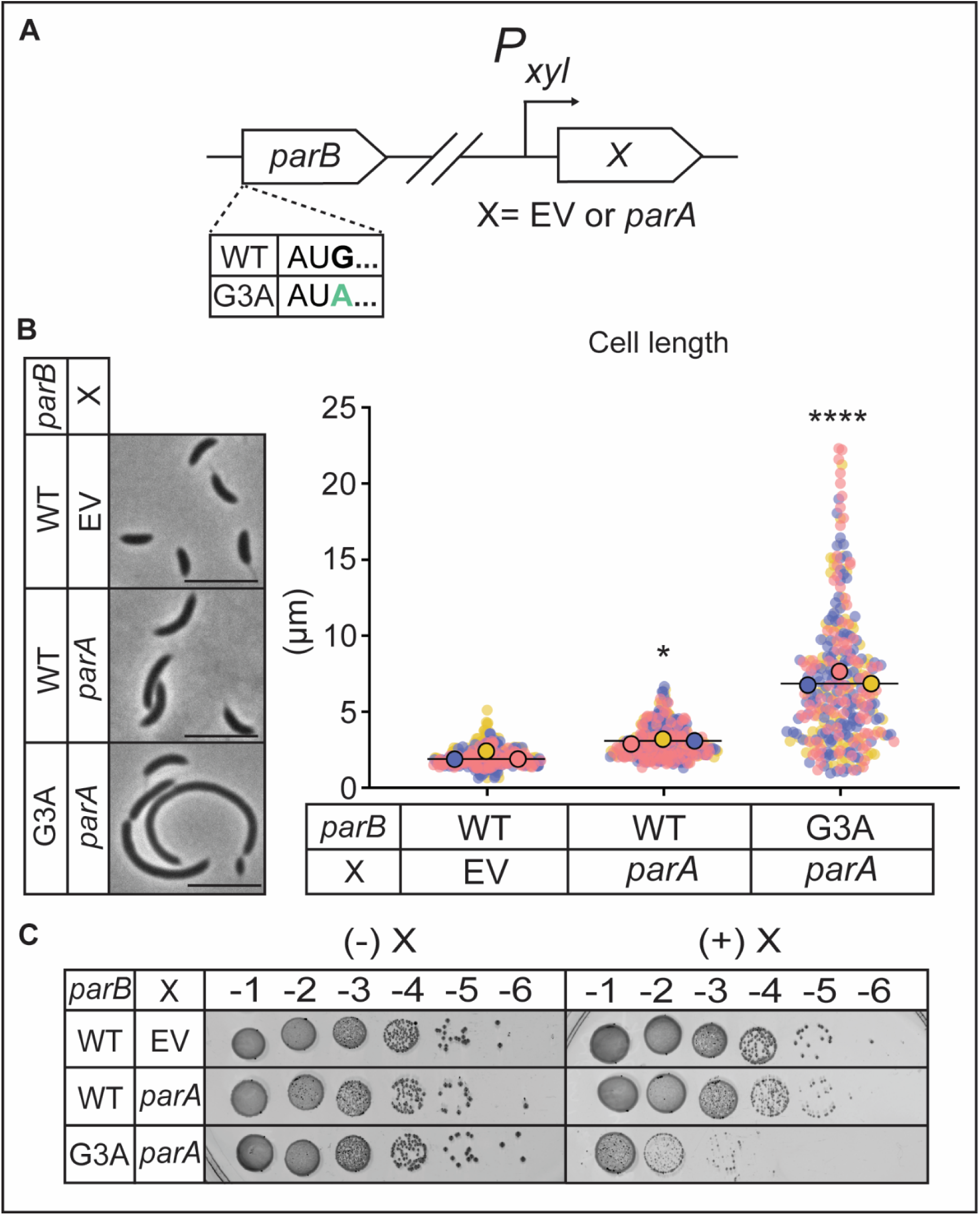
Δ10 ParB-only strains are highly susceptible to ParA fluctuations. **A.** Schematic of ParA inducible strains in a wild-type background or G3A background. **B.** (Left) Phase contrast images of ParA inducible strains in wildtype or G3A background after 6h of induction, grown in rich media (PYE) to exponential phase, scale 5 µm. (Right) Super plots showing cell length analysis of mixed population. Small points represent data points from three independent replicates, large dots represent median values (blue, pink, yellow). The horizontal lines represent the mean of three median values. One-way ANOVA was performed using population mean values (N=3) to compare values for each measurement. **C.** Colony-forming units showing viability of ParA inducible strains in WT or G3A background compared to empty vector (EV). G3A shows a viability defect compared to WT and EV.

### ParB’s N-terminus determines parS substrate specificity independent of the DNA-binding domain

To determine whether the loss of viability arose solely from reduced ParB levels or reflected an intrinsic functional defect in the Δ10 variant, we performed biochemical characterization of ParB *in vitro*. To directly compare DNA-binding properties, full-length and Δ10 ParB proteins were purified to homogeneity following heterologous expression in *Escherichia coli* BL21 (Methods). Electrophoretic mobility shift assays (EMSAs) revealed a striking alteration in DNA-binding behavior. Whereas full-length ParB bound the centromeric *parS* site with high affinity and exhibited minimal nonspecific DNA interactions, the Δ10 variant lost this selectivity and instead displayed promiscuous nonspecific DNA binding (Figure 7A). The observation that the N-terminus influences ParB substrate specificity was unexpected, given that *parS* substrate recognition has been thought to depend solely on the helix-turn-helix motif of the central DNA-binding domain [64].

**Figure 7.**
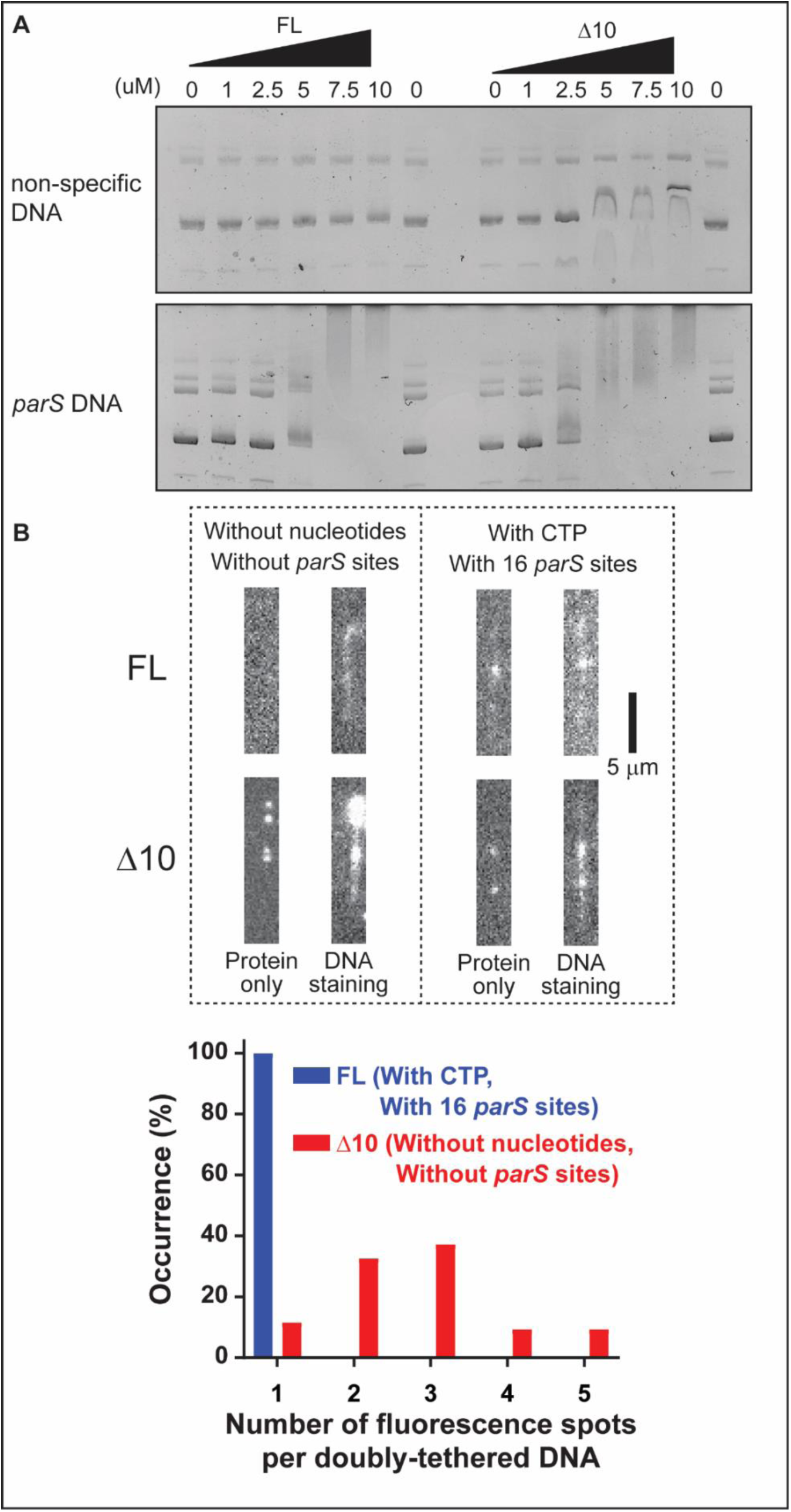
ParB’s N-terminus determines *parS* substrate specificity independent of the DNA-binding domain. **A.** EMSA showing FL and Δ10 binding to DNA using purified protein. FL and Δ10 ParB were incubated in the presence of (Upper) and empty vector pNPTS138 plasmid, or (lower) the *parS C. crescentus* sequence cloned into apNPTS138 plasmid. **B.** (Upper) Representative snapshot images of fluorescently labeled full-length and Δ10 ParB proteins. Either λ-DNA (devoid of any *parS* sequences) or engineered λ-DNA harboring 16 *parS* sites was used as a DNA substrate. After imaging solely protein, low concentrations of SYTOX Orange was introduced to stain the DNA. (Lower) Number of ParB fluorescence spots after washing unbound proteins by buffer flow.

To independently validate this loss of substrate selectivity, we examined the spatial distribution of ParB along DNA using direct visualization of ParB–DNA interactions. We employed a variant of single-molecule DNA flow-stretching assay due to its high sensitivity and versatility under a total internal reflection fluorescence (TIRF) microscope [65]. When full-length ParB was introduced to bacteriophage λ-DNA (without any *parS* sites) in the absence of CTP, more than 95% of doubly-tethered DNAs did not exhibit any ParB fluorescence signal (Figure 7B). However, when we performed the same experiment using the engineered 16-*parS* λ-DNA in the presence of CTP, bright fluorescent signals were detected, and the DNA staining by SYTOX Orange showed that the fluorescent signals were from DNA-bound full-length ParB proteins (Figure 7B). All the doubly-tethered DNAs (Supplementary Figure 2) had only one fluorescence focus per DNA (*n*=47), consistent with *C. crescentus* ParB accumulation on a closed *parS* DNA substrate in the presence of CTP [26]. Furthermore, the focus was found to be in the middle of the engineered 16-*parS* λ-DNA (where *parS* sites are incorporated), implying the specific ParB-*parS* interaction in the presence of *parS* and CTP. When the Δ10 ParB protein was tested with λ-DNA (without *parS* sites) in the absence of CTP, all the doubly-tethered DNAs exhibited multiple fluorescently labeled ParB signals (mean±s.d. = 2.72±0.98) (*n*=43) (Figure 7B), in stark contrast to the full-length ParB counterpart. The inclusion of *parS* sites and CTP did not alter the behavior of Δ10 ParB protein in that multiple ParB protein signals were detected even from the off-center of DNA (away from *parS* sites) (Figure 7B). Collectively, our *in vitro* EMSA and variant single-molecule DNA flow-stretching assays demonstrate that the first 10 amino acids are critical for conferring the specific ParB-*parS* interaction.

## DISCUSSION

In this study, we provide mechanistic evidence that chromosomal ParB levels are post-transcriptionally regulated. This regulation is mediated by a secondary structure at the 5′ end of *parB* coding DNA sequence (CDS) in *C. crescentus*. Using a series of single-base mutations, we demonstrate that disruption of this secondary structure reduces *parB* transcript and ParB protein levels. Using *parB* merodiploid strains as well as the native chromosomal locus, we demonstrate that the determinants of post-transcriptional regulation reside within the 5′ end and are largely independent of the surrounding sequence context. Together, these findings establish a previously unrecognized mechanism that fine-tunes ParB abundance to preserve the ParA–ParB balance.

In *C. crescentus*, the detection of post-transcriptional regulation was facilitated by the presence of two methionine codons that had previously been used indistinguishably as start sites. We propose that analogous regulation may be widespread in bacteria. Consistent with this, the *parB* genes of *Myxococcus xanthus* and *Streptococcus pneumoniae* each encode two potential start codons in close proximity (at amino acid positions 1 and 6), raising the possibility that a similar dual-start architecture underlies post-transcriptional tuning in these species. Furthermore, *parAB* operons associated with *gidAB* have been reported to contain a +1 transcriptional start site upstream of *parB* [66, 67], suggesting that independent regulation of *parB* expression is a more general feature of bacterial chromosome maintenance. Although the ∼30 bp sequence responsible for the regulatory secondary structure in *C. crescentus* does not appear to be broadly conserved, secondary structure-based regulation does not require sequence conservation but only the capacity to form a functionally equivalent structure. Identifying such structures in other species will require systematic structural probing rather than sequence comparison alone.

Another major finding of this study is that the first 10 amino acids of the ParB N-terminal domain contribute to substrate specificity. Using EMSA and single-molecule DNA flow-stretching assays, we demonstrate that full-length ParB binds *parS* with high affinity and selectivity, while Δ10 ParB loses this discrimination and instead binds DNA nonspecifically. This result was unexpected because the helix-turn-helix motif responsible for *parS* recognition resides in the central DNA-binding domain, and because *C. crescentus* ParB lacks the positively charged C-terminal domain residues that confer nonspecific DNA binding in other species [33, 44, 68–70]. Our single-molecule assays extend this finding by directly visualizing that full-length ParB localizes to a single focus at the *parS*-containing region of engineered λ-DNA in the presence of ParB’s co-substrate CTP, whereas Δ10 ParB distributes across multiple sites along the DNA regardless of *parS* or CTP. Previous studies have shown that specificity of ParB, and the related protein Noc, is largely encoded by residues at the protein–DNA interface within the central domain [64]. Our results expand this view by demonstrating that the N-terminal domain can serve as an additional determinant of substrate selectivity, independent of the canonical DNA-binding interface. Our findings add substrate selectivity to the growing list of functions attributed to the N-terminus of chromosomal ParB proteins, which include interaction with the partner protein ParA, CTP binding, higher-order complex assembly, and liquid–liquid phase separation [10, 24, 71, 72]. Together, these observations further underscore the functional importance of this domain.

ParB biochemical properties appear to be adapted to function in a species-specific manner. In *C. crescentus*, approximately ∼80% of ParB molecules are predicted to be localized at *parS* [20]. In contrast to *B. subtilis*, where near-equimolar concentrations of ParB and ParA are sufficient to stimulate robust ParA ATPase activity [11], *C. crescentus* ParB exhibits a relatively weak capacity to activate ParA [20]. Consequently, efficient stimulation of ParA ATPase activity in *C. crescentus* likely requires the assembly of higher-order ParB complexes at *parS*, resulting in locally elevated ParB concentrations [20]. Within this context, the loss of viability observed in G3A cells expressing the Δ10 ParB variant can be rationalized. The truncated N-terminal ParB binds DNA nonspecifically and fails to accumulate at sufficiently high levels at *parS*, thereby limiting its ability to stimulate ParA ATPase activity. Schnabel *et al*. have demonstrated the importance of ParB clamp closure in the presence of *parS* and CTP for efficient ParA-ParB interaction [45]. Our results show that the Δ10 construct is unresponsive to *parS* and CTP, which is another factor contributing to poor ParA ATPase stimulation. Therefore, in the presence of slightly elevated ParA levels, this imbalance likely leads to the persistence of ParA–ATP dimers, defective chromosome segregation, and ultimately loss of viability.

This study reveals that the 5′ end of the *C. crescentus parB* is a multifunctional regulatory element: it encodes a regulatory RNA secondary structure that controls ParB abundance and simultaneously specifies N-terminal residues required for *parS*-selective DNA binding. This dual functionality suggests that constraints on the 5′ end of *parB* extend beyond coding sequence requirements and reflect an integrated solution for coordinating both the quantity and activity of ParB. Given the conservation of ParABS-mediated chromosome segregation across bacteria, this dual-function architecture may reflect a general evolutionary solution for coupling ParB abundance and activity to maintain the precise stoichiometric balance required for faithful chromosome inheritance.

## MATERIAL AND METHODS

### Bacteria strain and growth conditions

Strain, plasmid, and oligonucleotide descriptions are listed in the Supplementary Table S3, S4, and S5. *C. crescentus* strains were grown in rich media (PYE) from freezer stocks at 30⁰C and 180 rpm. Exponentially growing cells were used for all experiments. Liquid media was supplemented with 5 μg/ml Kanamycin. PYE plates were supplemented with 25 μg/ml kanamycin. Where necessary, 0.1% xylose was added to induce the xylose promoter (P*_xyl_*). PYE medium was used during *C. crescentus* strain constructions.

### Plating CFUs for viability

*C. crescentus* strains were inoculated from freezer stocks, grown overnight in rich media (PYE) at 30⁰C and 180 rpm, and normalized to OD_600_ = 0.1. Cultures were then serially diluted 1:10 in fresh media in a 96-well plate. Eight microliters of each dilution were spotted onto PYE plates supplemented with 0.1% xylose where needed. Plates were grown at 30⁰C for 2 days, then imaged.

### Microscopy

*C. crescentus* cells (1 ul) were spotted on agarose pads (1% agarose in M2G), then phase contrast micrographs were taken using the Zeiss Axio Observer 2.1 inverted microscope with AxioCam 506 mono camera (objective: Plan-Apochromat 100×/1.40 Oil Ph3 M27 [WD = 0.17 mm]), and Zen lite software. Analyses of data were performed with blinded samples. Cell length was analyzed using ImageJ/FIJI with MicrobeJ software.

### Immunoblotting

To collect samples, a mixed population of *C. crescentus* cells was normalized to OD_600_ = 0.2, and 0.1% xylose was added where needed. Cells were pelleted and resuspended in 60 µl of Cracking buffer, then boiled for 10 min at 90°C before storage at –20°C for Western blotting. Samples were run on a 4-20% SDS-PAGE gel and transferred to a PVD membrane using iBlot2. The membrane was blocked for 1 h at room temperature with 1× TBST (10 mM Tris–HCl, pH 8.0, 150 mM NaCl, 0.1% Tween 20) with 5% non-fat milk. Blot was incubated overnight at 4⁰C with 1:10 000 diluted α-ParB, and α-ClpP where indicated. The membrane was washed 3x in TBST for 5 min each. The blot was probed with secondary antibody α-rabbit IgG peroxidase (Sigma-Aldrich) at a dilution of 1:10,000 at room temperature for 1 hour. The blot was washed 3x in TBST for 5 min. The blot was then developed using SuperSignal West Pico PLUS Chemiluminescent Substrate (Thermo Fisher Scientific) and imaged using the ChemiDoc-MP (Bio-Rad). The density of each band was quantified using ImageJ software.

### Protein purification for DNA binding assays

Protein purifications were performed as described (Lim et al., 2014) with modifications in the Mera Lab. In short, BL21 cells carrying pTEV-ParB or pTEV-Δ10ParB were grown in Luria-Bertani broth containing 50 μg/ml ampicillin to an OD_600_ of ∼0.6 before adding 1 mM isopropyl β-D-1-thiogalactopyranoside (IPTG) (GoldBio) and grown at 18°C overnight. Cells were pelleted at 6000 rpm for 10 minutes then stored at -80°C. The cell pellets were resuspended in Buffer A (50 mM HEPES/KOH (pH 7.5), 300 mM NaCl, 10 mM Imidazole) supplemented with lysozyme (0.5 mg/ml) and protease inhibitor (cOmplete protease inhibitor) for 30 min at 4°C. The resuspended pellets were then sonicated 3×, and centrifuged at 13000 rpm for 45 minutes at 4°C. The clarified lysate was injected into a HisTrap^TM^ FF crude 5 ml column (Cytiva Life Sciences) equilibrated in buffer A at 1 ml/min and washed with four column volumes (CV) of buffer A. Then, a gradient (0-100%) of buffer B (50 mM HEPES/KOH (pH 7.5), 300 mM NaCl, 500 mM Imidazole) was injected into the column at 1 ml/min. Fractions containing ParB-His6 were pooled and incubated with TEV protease (0.1 mg/ml) for 1 hour at room temperature. The sample was then dialyzed in Buffer A overnight at 4°C. The sample was injected into a HisTrap^TM^ FF crude 5 ml column equilibrated with buffer A at 1 ml/min. The column was washed with four CV a gradient (0-100%) of buffer B was injected into the column at 1 ml/min. Fractions containing ParB or Δ10ParB were concentrated using a Pierce^TM^ concentrator PES 10K MWCO. Pooled samples were subjected to dialysis in storage lysis buffer (50 mM HEPES/KOH (pH 7.5), 300 mM NaCl, 20% glycerol).

### Protein purification and fluorescent dye labeling for single-molecule assays

Rosetta2(DE3) pLysS cells transformed with a plasmid encoding either full-length or Δ10 ParB-His6 were grown at 37 °C. When the OD_600_ reached 0.4-0.6, the protein overexpression was induced by 0.5 mM IPTG. After 4 hours at 37 °C, the cells were harvested and resuspended in ParB lysis buffer (20 mM Tris, pH 8.0, 1 M NaCl, 50 mM imidazole, 5 mM 2-mercaptoethanol) in the presence of protease inhibitor cocktail (Roche) and 0.1 mM phenylmethylsulfonyl fluoride (PMSF). It was snap frozen and stored at -80 °C. The cells were thawed, supplemented with an additional 0.9 mM PMSF (total 1.0 mM). The addition of 0.25 mg/ml lysozyme, followed by sonication, lysed the cells. After two rounds of centrifugation (11,000 g for 30 minutes and 20,133 g for 30 minutes), the supernatant was incubated with Ni-NTA resin in the presence of apyrase and 5 mM MgCl_2_ for at least 1 hour. The Ni-NTA column was washed with ParB lysis buffer and ParB salt reduction buffer (20 mM Tris, pH 8.0, 350 mM NaCl, 50 mM imidazole, 5 mM MgCl_2_, and 5 mM 2-mercaptoethanol). The ParB protein was eluted using an elution buffer containing 250 mM imidazole. The elution fractions that contain the desired ParB protein were consolidated and dialyzed against ParB dialysis/storage buffer (20 mM tris, pH 8.0, 350 mM NaCl, 10 % glycerol, and 5 mM 2-mercaptoethanol). The protein concentration was measured using NanoDrop. Proteins were incubated with an excess of sulfo-Cyanine3 NHS ester dye (Lumiprobe) at 4 °C overnight. Labeled proteins were separated from free dye using Micro Bio-Spin P-30 gel columns (Bio-Rad). The concentrations of each labeled protein and Cyanine3 dye were measured three times using a Nanodrop, and the average values were used as the final concentrations.

### DNA binding assays

FL Δ10 ParB DNA binding was tested by incubating increasing amounts of protein in individual reaction tubes containing a master mix (20 mM HEPES/KOH [pH 7.5], 1 mM DTT, 1 mM MgCl_2_, 5 ng/ml pNPTS138 plasmid [5.4 kb]/ pNPTS138 – *parS* plasmid [6.5 kb]) to a final reaction volume of 10 μl per reaction tube. The reactions were incubated for 15 min at 37°C before being run on a 1.5% agarose gel on TBE buffer + 1 mM MgCl_2_. Then, gel was stained with 1 μg/ml Ethidium Bromide (Sigma Aldrich) before being imaged on a ChemiDoc-MP imaging system. Taylor *et al*. previously showed that the inclusion of magnesium ions is essential in detecting specific DNA binding with *Bacillus subtilis* ParB [69]. Having their findings in mind, our EMSA assays with *C. crescentus* ParB were performed in the presence of magnesium ions.

### DNA substrate preparations for single-molecule assay

Unmodified 48.5-kb bacteriophage λ-DNA (without any *parS* sequences) was purchased from New England Biolabs (N3013S). The engineered λ-DNA with 16 *parS* sites in the middle was extracted and purified from the λ^16*parS*^ lysogen strain (a gift from Xindan Wang’s lab) as described in our previous publication [33], with a modification. For this study, the DNA extraction procedure using phenol was introduced before performing phenol/chloroform/isoamyl alcohol (25:24:1)- and chloroform-based DNA extraction steps. The bacteriophage λ-DNA features 12-nucleotide overhangs at both ends. Biotinylated oligos (5’-AGGTCGCCGCCC/3BioTEG/-3’ and 5’-GGGCGGCGACCT/3BioTEG/-3’) complementary to the 12-nucleotide overhangs were used to biotinylate both ends of the λ-DNA, as detailed previously [73].

### Single-molecule assay with doubly-tethered DNA

The cover glass in a microfluidic sample chamber was coated with partially biotinylated polyethylene glycol (PEG) to minimize nonspecific adsorption of proteins and DNA. After incubation with 0.25 mg/ml neutravidin (in EBB+RA buffer: 10 mM Tris, pH 8.0, 150 mM NaCl, 10 mM MgCl_2_, and 0.2 mg/ml recombinant albumin) and subsequent washing to remove unbound neutravidin, doubly biotinylated DNA was introduced. The unbound excess DNA was washed away by flowing in imaging buffer (10 mM Tris, pH 7.5, 100 mM NaCl, 2.5 mM MgCl_2_). Fluorescently labeled ParB proteins, either with or without CTP, were introduced at a flow rate of 30 μl/min. After each experiment, DNA was stained with SYTOX Orange to visualize its entire length, and the flow was stopped to confirm whether the DNA was singly- or doubly-tethered. All experiments were performed on a total internal reflection fluorescence (TIRF) microscope equipped with an electron multiplying charge coupled device (EMCCD) camera and a 532 nm laser. The exposure time for recorded movies is 100 milliseconds, with images taken every 200 milliseconds, resulting in five frames per second.

### Reverse Transcription Quantitative PCR

*C. crescentus* cells were inoculated from freezer stocks, grown overnight in rich media (PYE) at 30⁰C and 180 rpm, and normalized to OD_600_ = 0.6. Cells were then pelleted, and RNA was harvested using the Thermo Scientific™ GeneJET RNA Purification Kit. Samples were treated with DNase (Invitrogen TURBO DNase) to remove genomic DNA. cDNA synthesis and RT-qPCR was performed using Luna universal one-step RT-qPCR kit. Final concentration of RNA was 10ng for each reaction. To quantify transcription ruvA (CCNA_03345) was used as an endogenous control, with the following primers: 5′-cgagtgaggaagccgtagag-3′ and 5′-gaccctgttgcacatcgag-3′.

## RESOURCE AVAILABILITY

All data are contained within the manuscript.

## ACKNOWLEDGEMENTS

We are grateful to Professor Xindan Wang and her lab at Indiana University for generously providing the *E. coli* strain containing 16 *parS*-lambda DNA. We thank the Carver Metabolomics and Proteomics Cores at the University of Illinois Urbana-Champaign for performing the mass spectrometry analysis. The work reported in this publication was supported by the National Science Foundation (NSF) Science and Technology Center for Quantitative Cell Biology under Award Number 2243257, and by the National Institute of General Medical Sciences of the National Institutes of Health (NIH) under Award Numbers R01GM133833 & R35GM158016 for P.E. Mera and R35GM143093 for H. Kim.

