## Supplemental Data for "A dual-function regulatory element couples ParB expression and DNA substrate specificity"

**FL and Δ10 overexpression phenotype is consistent in CFP-*parB* tagged strain.**

To differentiate between ParB made from the native promoter and ParB made from the Pxyl promoter, we used a strain with native *parB* fluorescently tagged, thereby increasing the molecular weight of the ParB expressed from the native promoter (CFP-ParB: 58.59 kDa vs. ParB: 32.81 kDa). To ensure that the FL and Δ10 overexpression phenotypes were not strain-specific, we transformed the same Pxyl-FL and Pxyl-Δ10 plasmids into the CFP-*parB* strain. We performed cell size analysis (Figure S1A) and colony-forming unit assays (Figure S1B) and observed the same phenotype to FL and Δ10 overexpression in WT cells.

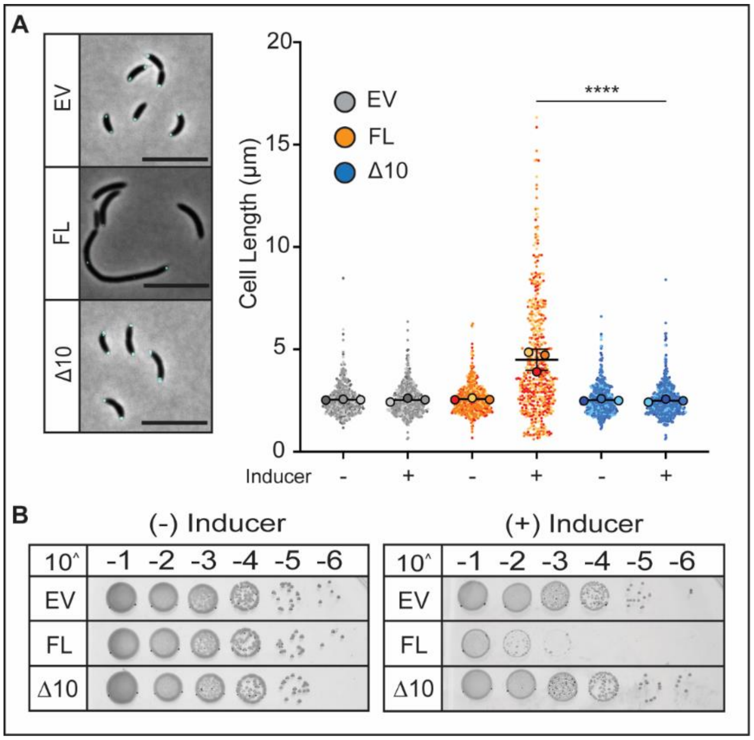


### Supplemental Figure 1. *C. crescentus* CFP-*parB* tagged strain displays the same phenotypes as WT when FL and Δ10 ParB are overexpressed.

**A.** (Left) Phase contrast images of FL and Δ10 inducible strains after 6h of 0.1% xylose induction, grown in rich media (PYE) to exponential phase, scale 5 µm. (Right) Cell length quantification of mixed population. Small points represent data points from three independent replicates, large dots represent median values. The horizontal lines represent the mean of three median values. One-way ANOVA was performed using population mean values (N=3) to compare values for each measurement. p(<0.0001) ****. The analyses were blinded n=~400 cells. **B.** Colony-forming units showing viability of inducible strains in comparison to empty vector (EV). EV and Δ10 show similar growth, while FL shows a viability defect.

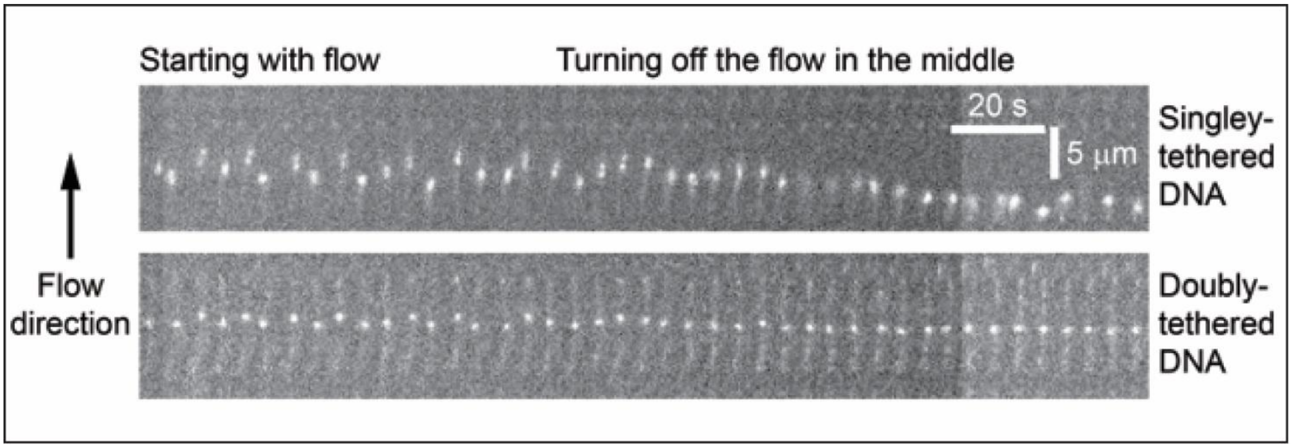


**Supplemental Figure 2. Distinction of doubly-tethered DNA from singly-tethered DNA.**

Whether only one end or both ends of DNA are tethered to the surface of a microfluidic sample chamber, the DNA is stretched by hydrodynamic flow. However, the removal of flow leads to the end-to-end length shortening for singly-tethered DNA, whereas the one for doubly-tethered DNA is unaffected by turning off the flow

Western blot patterns of ParB variants. For simplicity, we only showed the first time point and the last time point of each western blot in the main manuscript. For transparency, we display all of the time points for both accumulation and degradation of ParB (Figure S3A-D).

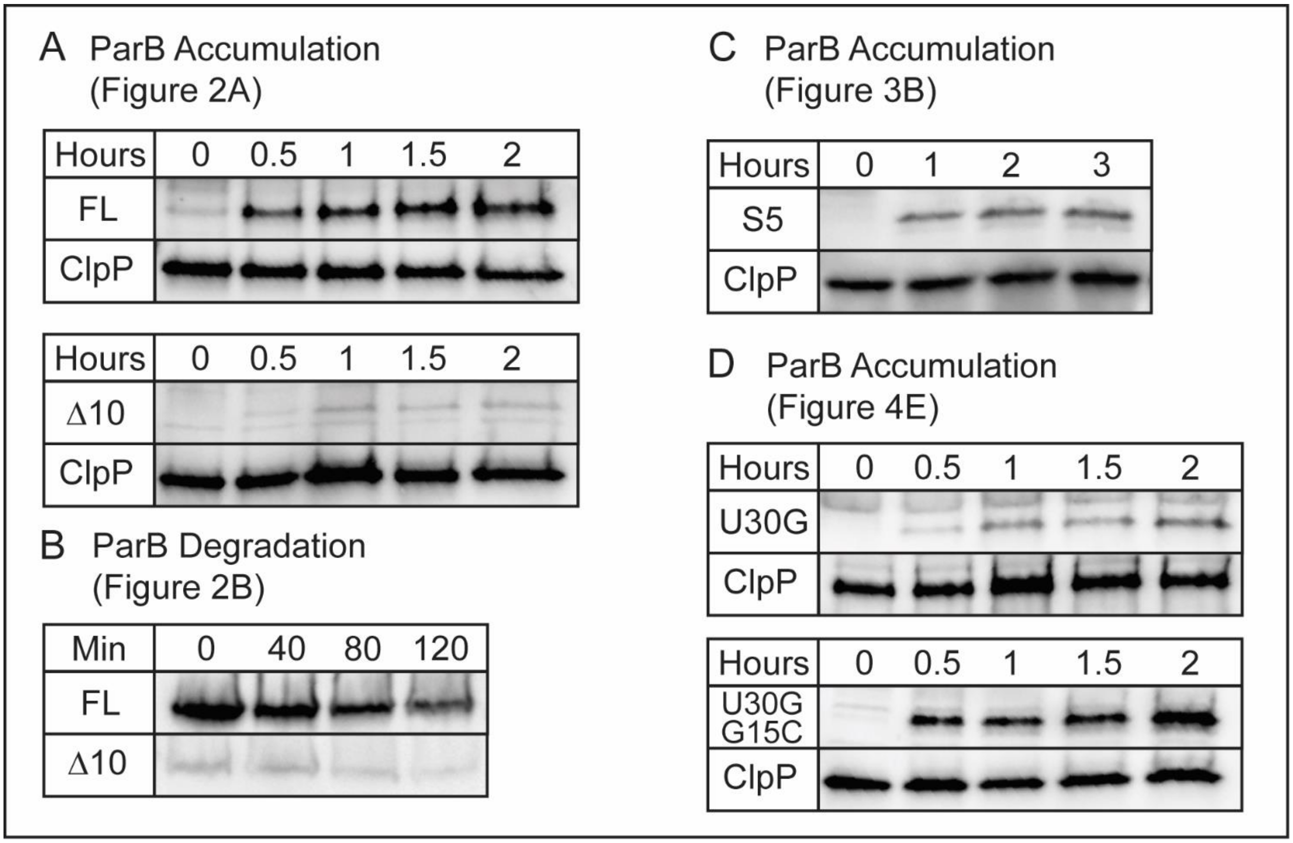


**Supplemental Figure 3. Whole western blot pictures**

Western blot accumulation and degradation over time of different ParBs. **A.** Western blot accumulation of FL and Δ10. ClpP is shown as a loading control. **B.** Western blot degradation of FL and Δ10. **C.** Western blot accumulation of S5. ClpP is shown as a loading control. **D.** Western blot accumulation of U30G and U30G + G15C. ClpP is shown as a loading control.

**Supplementary table 1:** Codon frequency table for *Caulobacter crescentus* CB15 [1]

| | [Triplet] | [Frequency per thousand] [number] | [Triplet] | [Frequency per thousand] [number] |
|---|---|---|---|---|
| Silent 5 | GAG | 38.5 [47647] | GAA | 15.5 [19152] |
| | GUC | 39.0 [48280] | GUG | 30.1 [37323] |
| | GUG | 30.1 [37323] | GUA | 1.2 [1517] |
| | GUG | 30.1 [37323] | GUU | 5.4 [6714] |
| | CCC | 18.7 [23146] | CCU | 3.5 [4338] |
| U30G | GGU | 9.1 [11281] | GGG | 11.3 [13983] |
| G15C | GUG | 30.1 [37323] | GUC | 39.0 [48280] |

**Supplementary Table 2**: List of strains used in this study

| Name | Relevant genotype | Reference |
|---|---|---|
| | CB15N (NA1000) | [2] |
| PM 566 | CB15N *parB::cfp-parB* Pxyl EV (kan$^R$) | [3] |
| PM 572 | CB15N *parB::cfp-parB* Pxyl -FL *parB* (kan$^R$) | This study |
| PM 644 | CB15N Pxyl – *parA* (kan$^R$) | This study |
| PM 652 | CB15N Pxyl -FL *parB* (kan$^R$) | This study |
| PM 702 | CB15N Pxyl -Δ10 *parB* (kan$^R$) | This study |
| PM 723 | CB15N *parB::cfp-parB* Pxyl Δ10 *parB* (kan$^R$) | This study |
| PM 732 | CB15N Pxyl EV (kan$^R$) | This study |
| PM 927 | CB15N *parB::*G3A *parB* | This study |
| PM 1118 | CB15N *parB::*G3A *parB* Pxyl - *parA* (kan$^R$) | This study |
| PM 1533 | CB15N Pxyl -S5 *parB* (kan$^R$) | This study |
| PM 1534 | CB15N *parB::cfp-parB* Pxyl -S5 *parB* (kan$^R$) | This study |
| PM 1738 | CB15N Pxyl -U30G *parB* (kan$^R$) | This study |
| PM 1739 | CB15N *parB::cfp-parB* Pxyl -U30G *parB* (kan$^R$) | This study |
| PM 1740 | CB15N Pxyl -U30G+G15C *parB* (kan$^R$) | This study |
| PM 1741 | CB15N *parB::cfp-parB* Pxyl -U30G+G15C *parB* (kan$^R$) | This study |

**Supplementary Table 3:** List of plasmids used in this study

| Plasmid name | Relevant genotype | Reference |
|---|---|---|
| pNPTS138 | Nonreplicating vector for allelic replacement (kanR or chlorR) oriT sacB | Alley M. R. K., unpublished |
| pXCHYC-2 | Integrating constructs encoding C-terminal mCherry fusions under the control of native PxylX (kan$^{R}$) | [4] |
| pDNA 69 | *parB*-(Δ10) cloned into pTEV-5 (amp$^{R}$) | This study |
| pDNA 245 | *parA* cloned into pXCHYC-2 (kan$^{R}$) | [3] |
| pDNA 264 | *mCherry* tag excised from pXCHYC-2 (kan$^{R}$) | [3] |
| pDNA 297 | *parB*-(FL) cloned into pXCHYC-2 (kan$^{R}$) | This study |
| pDNA 317 | *parB*-(Δ10) cloned into pXCHYC-2 (kan$^{R}$) | This study |
| pDNA 369 | pNPTS138 derivative to replace *parB* by *parB*(G3A) under the native $P_{parB}$ (kan$^{R}$) | This study |
| pDNA 578 | *parB*-(S5) cloned into pXCHYC-2 (kan$^{R}$) | This study |
| pDNA 609 | *parB*-(U30G) cloned into pXCHYC-2 (kan$^{R}$) | This study |
| pDNA 610 | *parB*-(U30G+G15C) cloned into pXCHYC-2 (kan$^{R}$)0 | This study |
| pDNA 626 | *parB*-(FL) cloned into pTEV-5 (amp$^{R}$) | This study |
| m0146 | *parB*-(FL) with a His-tag at the C-terminus (kan$^{R}$) | This study |
| m0148 | *parB*-(Δ10) with a His-tag at the C-terminus (kan$^{R}$) | This study |

**Supplementary Table 4:** List of primer oligonucleotides used in this study

| Plasmid name | Oligonucleotide sequence (5’→3’) | Use |
|---|---|---|
| pDNA 69 | Fwd - AAAAAACATATGTCCGAAGGGCGTCGTGGTCTG | *parB* (Δ10) cloned into pTEV-5 |
| | Rev - AAAAAAGCGGCCGCATCAGATCCCGCGCGTCAGTC | |
| pDNA 245 | Fwd - AAAGAGCTCTGGCGGCCTTGGCCTG | *parA* cloned into pXCHYC-2 |
| | Rev - AAAGCTAGCTTAGGCGGCCTTGGCCTG | |
| pDNA 264 | Fwd -TCGAGTTTTGGGGAGACGACCATATGTGCAGC CCGGGGGATCC | mCherry tag excised from pXCHYC-2 via Gibson assembly |
| | Rev - GTGCTGCAAGGCGATTAAG | |
| pDNA 297 | Fwd - AAACATATGGAGTCCGTCGTGGTGGGAGAG | *parB* (FL) cloned into pXCHYC-2 |
| | Rev - GCTAGCTCAGATCCCGCGCGTCAGTC | |
| pDNA 317 | Fwd - AAAAAACATATGTCCGAAGGGCGTCGTGGTCTG | *parB* (Δ10) cloned into pXCHYC-2 |
| | Rev - AAAGCTAGCTCAGATCCCGCGCGTCAGTC | |
| pDNA 369 | UP 500 bp Fwd - CGGCCGAAGCTAGCGAATTCGTGGAGCCCTGGAGGCGATCCG | pNPTS138 derivative to replace *parB* by *parB* (G3A) under the native $P_{parB}$ via Gibson assembly |
| | G3A Fwd - GTAGCTTGACTGAGATTGTGATAGAGTCCGTCGTGGTGGG | |
| | G3A Rev - CCCACCACGACGGACTCTATCACAATCTCAGTCAAGCTAC | |
| | DWN 500 bp Rev - CAAGCTTCTCTGCAGGATATCTGGATGGGCGATGTTCTCCTG | |
| pDNA 578 | Fwd – CAGACGCTCGAGTTTTGGGGAGACGACCATATGGAATCCGTG GTAGTTGGAGAGCCAGGT | *parB* (S5) cloned into pXCHYC-2 via site-directed mutagenesis |
| | Rev – TCGACCCAGACCACGACGCCCTTCGGACATACCTGGCTCTC CAACTACCACGGATTCCAT | |
| pDNA 609 | Fwd – CTCGAGTTTTGGGGAGACGACCATATGGAGTCCGTCG TGGTGGGAGAGCCCGGGATGTCC | *parB* (U30G) cloned into pXCHYC-2 via site-directed mutagenesis |
| | Rev – GGACATCCCGGGCTCTCCCACCACGACGGACTCCATA TGGTCGTCTCCCCAAAACTCGAG | |
| pDNA 610 | Fwd – CTCGAGTTTTGGGGAGACGACCATATGGAGTCCGTCGT CGTGGGAGAGCCCGGGATGTCC | *parB* (U30G+G15C) cloned into pXCHYC-2 via site-directed mutagenesis |
| | Rev – GGACATCCCGGGCTCTCCCACGACGACGGACTCCATA TGGTCGTCTCCCCAAAACTCGAG | |

| | | |
|---|---|---|
| pDNA 626 | Fwd - AAACATATGGAGTCCGTCGTGGTGGGAGAG<br>Rev - AAAAAAGCGGCCGCATCAGATCCCGCGCGTCAGTC | *parB* (FL) cloned into pTEV-5 |
| Lambda-BL1 Biotin | AGGTCGCCGCCC | Lambda DNA tagging with a biotin at one end |
| Lambda-BL2 Biotin | GGGCGGCGACCT | Lambda DNA tagging with a biotin at the other end |
